# Loop Assembly: a simple and open system for recursive fabrication of DNA circuits

**DOI:** 10.1101/247593

**Authors:** Bernardo Pollak, Ariel Cerda, Mihails Delmans, Simón Álamos, Tomás Moyano, Anthony West, Rodrigo A. Gutiérrez, Nicola Patron, Fernán Federici, Jim Haseloff

**Affiliations:** Department of Plant Sciences, University of Cambridge, Cambridge CB2 3EA, United Kingdom.; Departamento de Genética Molecular y Microbiología, Facultad de Ciencias Biológicas, Pontificia Universidad Católica de Chile, Santiago, Chile, Fondo de Desarrollo de Áreas Prioritarias, Center for Genome Regulation, Millennium Institute for Integrative Systems and Synthetic Biology, Pontificia Universidad Católica de Chile, 8331150, Santiago, Chile.; Department of Plant and Microbial Biology, University of California, Berkeley, California, USA.; Earlham Institute, Norwich Research Park, Norwich NR4 7UZ, United Kingdom.

**Keywords:** Standardised DNA assembly, Type IIS assembly, UNS, recursive DNA assembly, *LoopDesigner*, Open-MTA, Loop assembly, combinatorial assembly, synthetic promoters, common syntax, DNA fabrication

## Abstract

High efficiency methods for DNA assembly are based on sequence overlap between fragments or Type IIS restriction endonuclease cleavage and ligation. These have enabled routine assembly of synthetic DNAs of increased size and complexity. However, these techniques require customisation, elaborate vector sets and serial manipulations for the different stages of assembly. We present Loop assembly, based on a recursive approach to DNA fabrication. Alternate use of two Type IIS restriction endonucleases and corresponding vector sets allows efficient and parallel assembly of large DNA circuits. Plasmids containing standard Level 0 parts can be assembled into circuits containing 1, 4, 16 or more genes by looping between the two vector sets. The vectors also contain modular sites for hybrid assembly using sequence overlap methods. Loop assembly provides a simple generalised solution for DNA construction with standardised parts. The cloning system is provided under an OpenMTA license for unrestricted sharing and open access.

## Introduction

Standardised approaches to the assembly of large DNAs have played an important role in the development of systematic strategies for reprogramming biological systems. This began with the implementation of idempotent assembly methods based on DNA digestion/ligation using standardised nested restriction endonuclease (RE) sites, such as the BioBrick assembly method^1, 2^. More recently, assembly techniques that enabled the parallel assembly of multiple components in a single reaction have been established. These include methods that utilise long-sequence overlaps^3–10^, systems reliant on *in vivo* recombination^11–13^, and Golden Gate^14^ based methods that rely on selective digestion and re-ligation of plasmid DNAs with Type IIS RE^15–21^. Type IIS and long-overlap based methods have allowed increased scale and efficiency of DNA circuit assembly, while *in vivo* recombination remains the method of choice for genome-scale manipulations^12, 22–26^.

Gibson assembly, a sequence overlap-based method, was developed for the synthesis and assembly of *Mycoplasma* genomes^25, 26^ and enabled assembly of DNAs up to several hundred kb in one-pot isothermal reactions^3^. This method has been widely adopted by the synthetic biology community, being scar-free, versatile and relatively efficient. However, Gibson assembly generally relies on the use of oligonucleotides to perform *in vitro* amplification of DNA fragments, which can be error-prone. The method is also sensitive to sequence composition and repeats, and can be unreliable for inexperienced users. Efforts have been made to standardise and streamline Gibson assembly by including flanking unique nucleotide sequences (UNS) that can be used as long overlaps for cloning of transcription units (TUs) into larger constructs^27^. Gibson assembly can work efficiently for assembling a small number of TUs but relies increasingly on complementary methods for larger constructs. Perhaps due to the flexible nature of Gibson assembly, a standard for composing elemental parts into TUs has not been proposed yet. Thus, laboratories that employ Gibson assembly rely on their own set of rules and templates for DNA parts, and there has been no community-wide effort to develop a common standard.

In contrast, Type IIS assembly systems are virtually free of *ad hoc* design, and are highly efficient^28^. The methods do not require PCR amplification or fragment isolation, and allow parallel assembly of a large number of DNA parts. They rely on Type IIS RE to generate fragments with defined compatibility, and to recut unwanted side products of ligation in a one-pot system. While this approach can be ‘scarless’, the application of standard overhangs (fusion sites) for DNA parts with a defined function (e.g. promoter, CDS, terminators)^14^ allows the same DNA parts to be re-assembled into multiple constructs without redesign or modification. This allows efficient and robust parallel assembly of multiple parts in reactions that are simple to automate. Therefore, a common syntax has been proposed by developers and adopters of Type IIS cloning methods. This standard defines an unambiguous arrangement of 12 Type IIS overhangs that form boundaries between functional domains found within a generalised eukaryote gene^29^. The common syntax is based on the widely used MoClo and GoldenBraid standards, and has found acceptance in the plant field, where compatible parts are termed PhytoBricks. This ensures that these Type IIS assembly systems, which rely on BsaI as an entry point, can share a common stock of standardised DNA parts to be shared and used in an off-the-shelf manner. The establishment of a common standard for stock DNA parts also provides a prevailing syntax that enhances transferability and reproducibility for compiling genetic instructions in different labs. Assembly of an exact copy of a genetic construct is possible simply by knowing its composition, eliminating unnecessary *ad hoc* design and enabling simple abstract descriptions that contain a precise implied sequence. However, Type IIS assembly systems require the refactoring or ‘domestication’ of DNA parts. Domestication refers to the elimination of RE sites present in the DNA sequence prior to its use in the assembly system. To date, the most frequently used REs have been BsaI, BsmBI and BpiI, which have 6 bp recognition sites that, while not frequent, are regularly encountered in eukaryotic genomes. Type IIS REs such as SapI and AarI with 7 bp recognition sites can be used to lower the chance of finding illegal sites, and are used in the Electra^TM^ (ATUM) and GeneArt^TM^ (ThermoFisher) kits, respectively. Type IIS based systems have found rapid acceptance in the synthetic biology field due to the need for robustness, scalability and compatibility with automated assembly methods. Since synthetic biology is already at the point where constructs can consist of multiple logic gates^30^, entire biosynthetic pathways^31^ or engineered genomic DNA^32^, robust assembly methods such as Type IIS assembly are essential to enable fabrication of higher-order genetic constructs.

Despite much progress in the technical aspects of DNA construction and part reusability, restrictive intellectual property (IP) practices and material transfer agreements (MTA) can hinder the sharing of DNA components in both the public and private sectors, delaying experimental work through paperwork and legal consultation. For this purpose, an international effort is underway to establish the OpenMTA (http://www.openmta.org) as a way of expediting the sharing of biological materials. The OpenMTA provides a legal template for free and unrestricted distribution of materials, providing a formal mechanism for effectively placing materials in the public domain, in a way that extends existing practices. Open sharing of DNA assembly systems and parts through the OpenMTA will facilitate the engineering of new solutions for problems in human health, agriculture and the environment, such as those identified as Sustainable Development Goals by the United Nations (http://www.un.org/sustainabledevelopment) and Global Grand Challenges by the Gates Foundation (https://gcgh.grandchallenges.org).

Here we present Loop assembly, a versatile, simple and efficient DNA fabrication system based on recursive DNA assembly. In our method, Type IIS assemblies are performed through iterated ‘loops’. Two sets of plasmid vectors are provided, which allow alternating assembly cycles. First, Level 0 parts, defined by the PhytoBrick common syntax, are assembled into Level 1 transcription units in each of four odd-numbered vectors using BsaI. Second, four Level 1 modules can then be assembled into a Level 2 construct in each of the four even-numbered vectors using SapI. Following this, Level 2 constructs can be combined by cloning back into odd-numbered vectors, using BsaI, to create Level 3 assemblies containing up to 16 transcription units each (Fig. 1). The iterative process of combining genetic modules, four at a time, can be continued without theoretical limit, alternating assembly steps between odd and even Loop vectors. Since levels are used recursively, it is possible to create hybrid levels that can contain a mixture of parts from different levels of the same parity (i.e. Level 2 vectors combined with elements from Level 0 vectors). As well as Type IIS assembly, the system integrates long-overlap assembly methods. In this way, four TUs can be assembled into multiple transcription units by using alternative methods such as Gibson Assembly via flanking UNS^27^. In addition, we have developed *LoopDesigner*, a software framework for *in silico* sequence handling and assembly design. The software tools are open source and available through *Github*, and Loop assembly vectors are provided through the OpenMTA for unrestricted use. We have developed and tested the Loop assembly system in different laboratories and provide data to support the efficiency and robustness of the method. We have assembled over 200 constructs with up to 16 TUs and over 38 kb in size. We have tested Loop constructs *in planta* and validated their function in transgenic *Marchantia polymorpha,* and through transient expression in *Arabidopsis thaliana* protoplasts.

## Results

### Loop assembly

Loop assembly consists of 2 sets of plasmids that participate in a cyclic assembly process. Type IIS restriction endonucleases BsaI and SapI are used alternately for recursive assembly of genetic modules into a quartet of odd (L1, L3,…) and even (L2, L4,…) receiver plasmids. At each step in the assembly ‘loop’, 4 genetic modules are combined into a receiver plasmid (Fig. 1a). Odd and even-level plasmids use alternating types of antibiotic selection, kanamycin resistance for odd-levels (pOdd plasmids) and spectinomycin resistance for even-levels (pEven plasmids), to enable the use of one-pot digestion-ligation assembly reaction^14^. Loop assembly allows parallel combinatorial assembly into receiver plasmids, permitting rapid and flexible construction of genetic libraries with multiple TU arrangements. At each level (except for TU assembly from L0 parts), four parental plasmids are required, leading to an exponential increase in the number of TUs by a factor of 4 per level (Fig. 1b, Supplementary Fig. 1).

**Figure 1.**
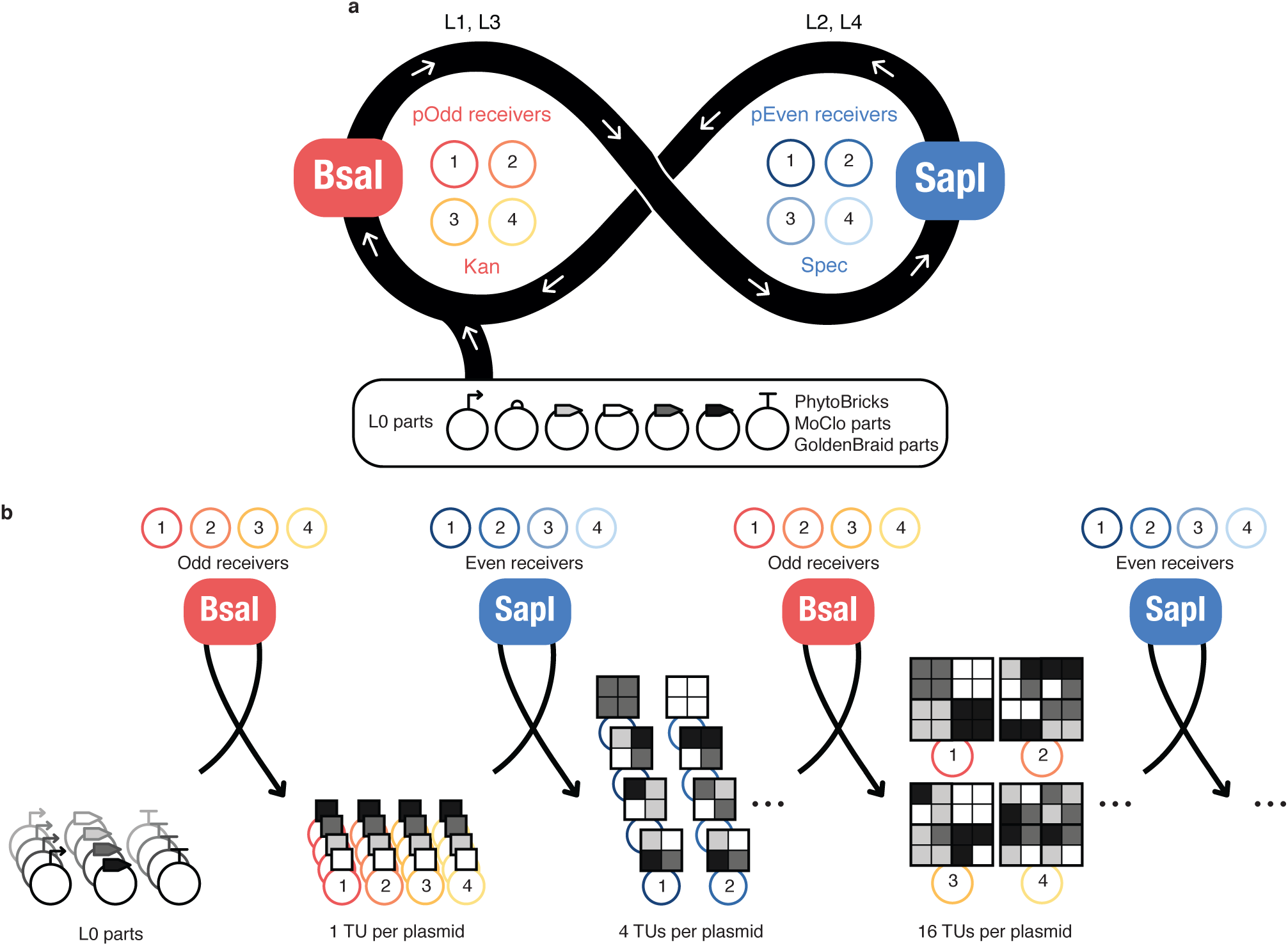
Overview of Loop assembly. a, Loop assembly workflow. L0 parts are assembled to L1 TUs into one pOdd receiver by BsaI-mediated Type IIS assembly. L1 TUs are assembled to L2 multi- TUs into one pEven receiver by SapI-mediated Type IIS assembly. This workflow is then repeated for higher-level assemblies. Only four odd level and four even level receiver plasmids are required for Loop assembly. **b, Combinatorial and exponential assembly**. L0 parts can be assembled to L1 TUs into any of the four positions of odd receivers. Genetic modules can be easily be swapped in each TU arrangement and receiver position. L1 TUs can then be assembled into L2 multi-TUs with variable combinations of the L1 TUs, also into any of the four positions of the even receivers. Each round of assembly generates four assembled plasmids and consequent rounds of assembly increase the number of TUs by a factor of four, leading to an exponential increase in TU number.

Odd numbered Loop acceptors contain a pair of divergent BsaI sites that are removed in the cloning reaction. These are flanked by a pair of convergent SapI sites for assembly into even numbered plasmids. In contrast, even numbered plasmids contain a pair of divergent SapI sites flanked by convergent BsaI sites (Fig. 2a). Upon digestion, plasmids release DNA fragments with specific overhangs that define the direction and position in the assembly. The overhangs created by the digestion of odd numbered Loop acceptors by BsaI allow the construction of transcription units from any parts that are compatible with the PhytoBrick standard^29^ (such as MoClo and GoldenBraid L0 parts), if free of SapI sites. BsaI overhang sequences are termed A, B, C, E and F, with A and F designated as flanking acceptor-overhangs, and SapI overhang sequences are termed α, β, γ, ε and ω, with α and ω designated as flanking acceptor-overhangs. Examples of odd and even-level assemblies for receiver plasmids are shown in (Fig. 2bc).

**Figure 2.**
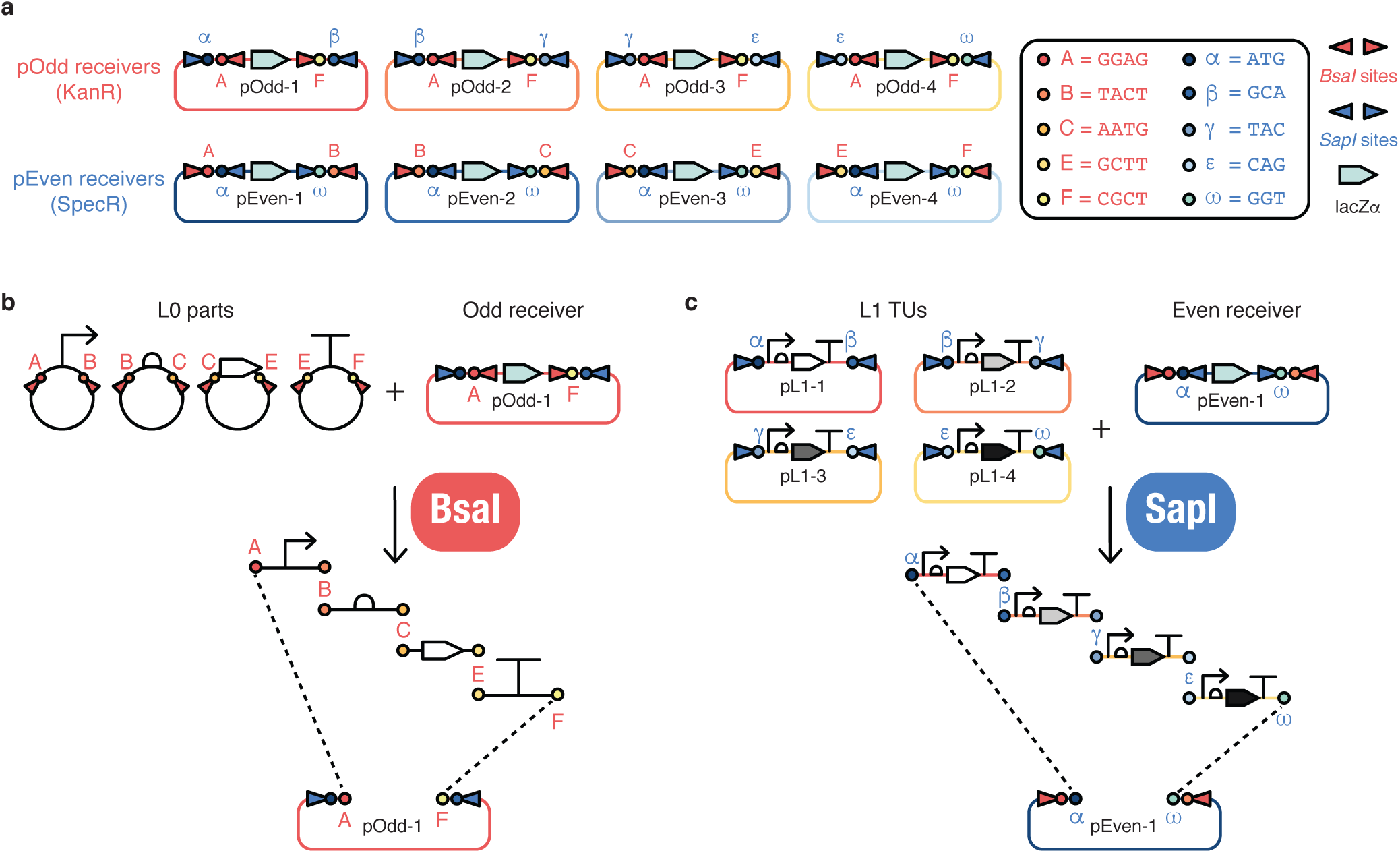
Loop assembly schema. a, Loop receiver plasmids. Each of the four pOdd and pEven receivers plasmids has a specific set of SapI (3 bp) and BsaI (4 bp) convergent overhangs respectively, required for higher level assembly. Odd receivers contain diverging BsaI restriction sites and acceptor-overhangs according to the common syntax, making them compatible for cloning L0 parts into pOdd plasmids. They contain SapI converging sites with donor-overhangs for directing SapI-mediated Type IIS assembly into even-level receivers. pEven plasmids have SapI diverging restriction sites and acceptor-overhangs to receive parts from pOdd plasmids. For higher-level assemblies, pEven plasmids contain converging BsaI sites with donor-overhangs for BsaI-mediated Type IIS assembly into pOdd plasmids. **b, Loop odd-level assembly**. L0 DNA parts containing overhangs defined in the common syntax are assembled into a Loop odd-level receiver. BsaI digestion releases the DNA modules, which are assembled into an even-level receiver by directional assembly defined by 4 bp overhangs. pOdd plasmids contain A and F overhangs as acceptor-overhangs for receiving parts, which are flanked by convergent SapI restriction sites with 3 bp donor-overhangs for further assembly. **c, Loop even-level assembly**. Four previously assembled pL1 TUs are assembled into a pEven plasmid. SapI digestion releases TUs from pL1 plasmids, which are assembled into an even-level receiver by directional assembly defined by 3 bp overhangs. pEven plasmids contain α and ω overhangs as acceptor-overhangs, which are flanked by convergent BsaI restriction sites with donor-overhangs defined in the common syntax required for further assembly.

Each reaction requires four donor plasmids (or spacers) for successful assembly into a receiver of the next level. In order to provide a replacement for missing modules in assemblies, we designed ‘universal spacer’ parts comprised of 200 bp of random DNA sequence without BsaI and SapI sites. Universal spacer plasmids are provided for odd (pOdd-spacer) and even levels (pEven-spacer) and contain flanking sites that allow them to be assembled directly into any of the four receiver plasmids of their intended level, to then be used for filling positions in assemblies (**Supplementary Fig. 2**).

### Assembly of synthetic promoters

The recursive nature of Loop assembly allows the mixing of parts from different odd or even levels. For example, a multimeric promoter might be constructed from elemental parts through recursive assembly. Figure 3 shows the generation of synthetic promoters by cloning L0 functional domains (e.g. TF recognition sites and minimum promoter sequences) into specific L1 plasmid positions, which determine the order of arrangement in the next L2 assembly. Different TF recognition sites can be used in positions 1 (α and β overhangs), 2 (β and γ overhangs) and 3 (γ and ε overhangs), while a minimal promoter sequence is placed in position 4 (ε and ω overhangs) of L1 receiver plasmids. These elements can then be composed in specific order. In this example, different combinations of TF binding sites and minimal promoter were cloned into positions 1 (A and B overhangs) and 2 (B and C overhangs) of L2 receiver plasmids. The resulting composite promoter elements could be mixed with standard L0 gene parts, to create a customised hybrid gene assembly in a odd-level plasmid (Fig. 3a).

Using this approach, we assembled 3 fluorescent reporters with synthetic promoters comprised of multimeric binding sites. The promoters included multimeric binding domains for the transcription factors GAL4^33–35^ (a 47 bp dimeric binding domain, 2xUAS_GAL4_), and HAP1^36^ (a 49 bp dimeric binding domain, 2xUAS_HAP1_), a cytokinin operator (a 90 bp type-B ARABIDOPSIS RESPONSE REGULATOR binding domain^37^, CK_OP_) and a minimal CaMV 35S promoter^38^ (min35S) derivative (Federici and Haseloff, unpublished results). Resulting reporters were composed of the same elements but with differing arrangements. Each reporter contained 3 dimeric binding domains for GAL4, 3 dimeric binding domains for HAP1, one dimeric CK operator binding domain and the minimal CaMV 35S promoter. Binding domains for each TF were placed together in sets and each composite promoter represented a specific permutation of the positioning of the binding domains (see **Supplementary Text 1)**. The composite synthetic promoters, which were the result of 20 different assembly reactions, were verified through sequencing and showed no sequence errors.

The recursive nature of Loop assembly also enables hybrid assemblies of multiple TUs derived from parental plasmids from different levels (i.e three Level 1 and one Level 3 plasmids). These can be assembled into a hybrid even receiver plasmid, providing further flexibility in the fabrication of genetic constructs (Fig. 3b).

**Figure 3.**
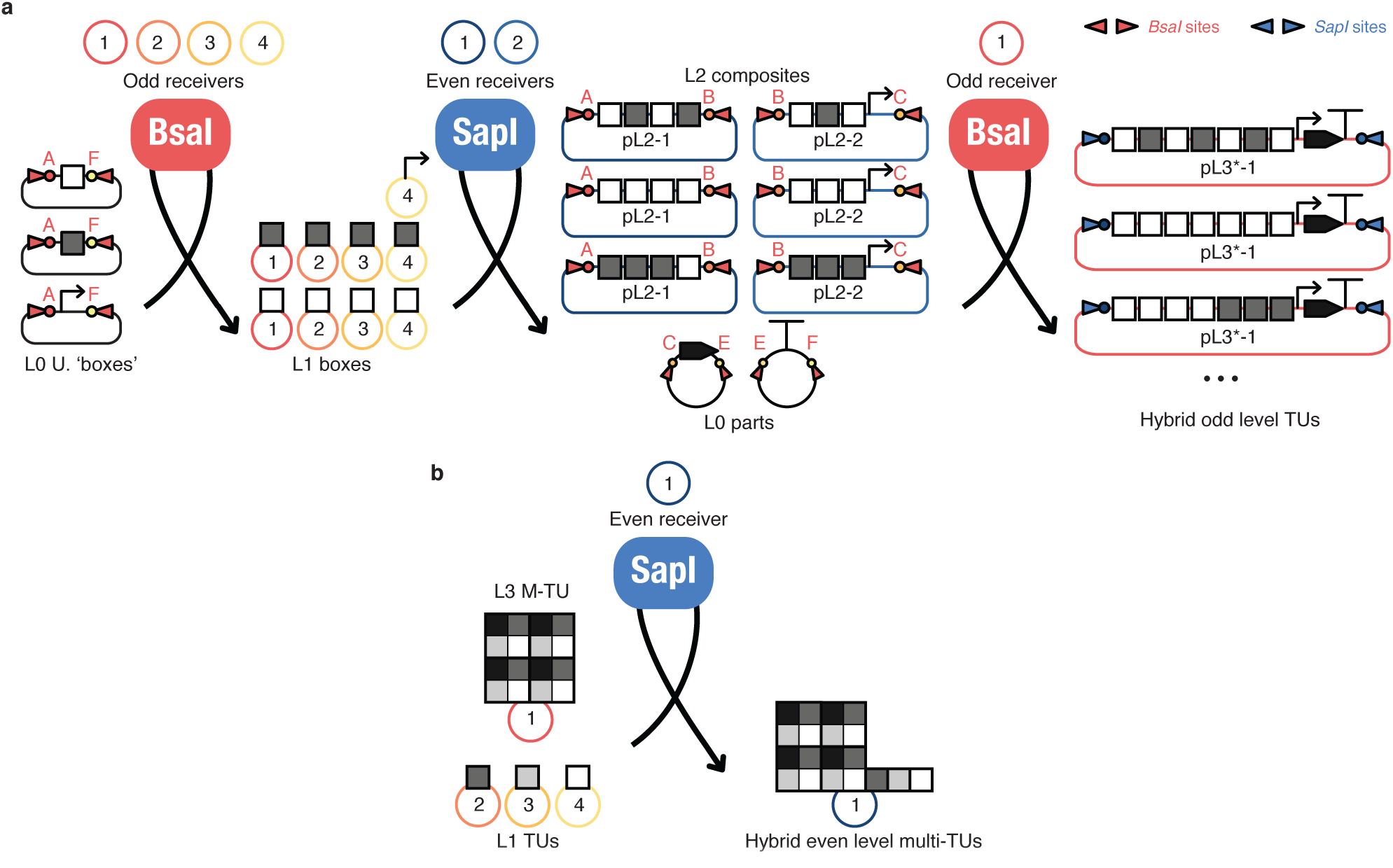
Hybrid assembly. a, Synthetic promoter assembly. L0 universal (U.) ‘boxes’ are assembled into odd level receivers into any given position. L1 boxes are then assembled into L2 composites with differing arrangements into positions 1 and 2. L2 composites in positions 1 and 2 are used in a hybrid assembly with L0 parts to generate a hybrid odd level TU with a synthetic promoter composed of the L0 boxes in the defined arrangement. **b, Mixed level assembly**. L3 and L1 parts are assembled into a even level receiver generating a hybrid even level multi-TU plasmid.

### UNS for standardised overlap assembly

We have added features to the Loop assembly vectors to enable the combination of Type IIS and standardised long overlap assembly techniques. Loop plasmids contain unique nucleotide sequences^27^ (UNS) sites that allow the use of standard primers for the amplification of TUs. The UNS allow facile PCR amplification of TUs derived from Type IIS DNA parts (PhytoBricks, MoClo and GoldenBraid), since these can be assembled into UNS-flanked TUs by BsaI-mediated Type IIS assembly. Alternatively, TUs can be assembled from PCR-fragments or DNA synthesis into Loop plasmids by overlap assembly methods such as Gibson assembly (Fig. 4a). Each Loop plasmid contains two flanking UNS and a terminal UNS_x_. TUs can be assembled into a multi-TU destination plasmid (pUNSDest) by using overlap assembly methods (Fig. 4b). UNS have been designed following a number of guidelines to provide enhanced performance in PCR reactions and overlap assembly. Design rules are listed in **Supplementary Text 2** and sequences provided in **Supplementary Table 1**. Forward and reverse standard primers correspond to the first 20 bp of each UNS in both forward and reverse complement orientations, respectively, and are provided in **Supplementary Table 2**. UNS have the advantage that they are designed for highly efficient PCR with standard conditions (60 ºC, 35 cycles), resulting in single amplicons with high yields (**Supplementary Fig. 3**). This eliminates the need for gel purification during the workflow of Gibson assembly, if appropriate on-column purification is performed.

**Figure 4.**
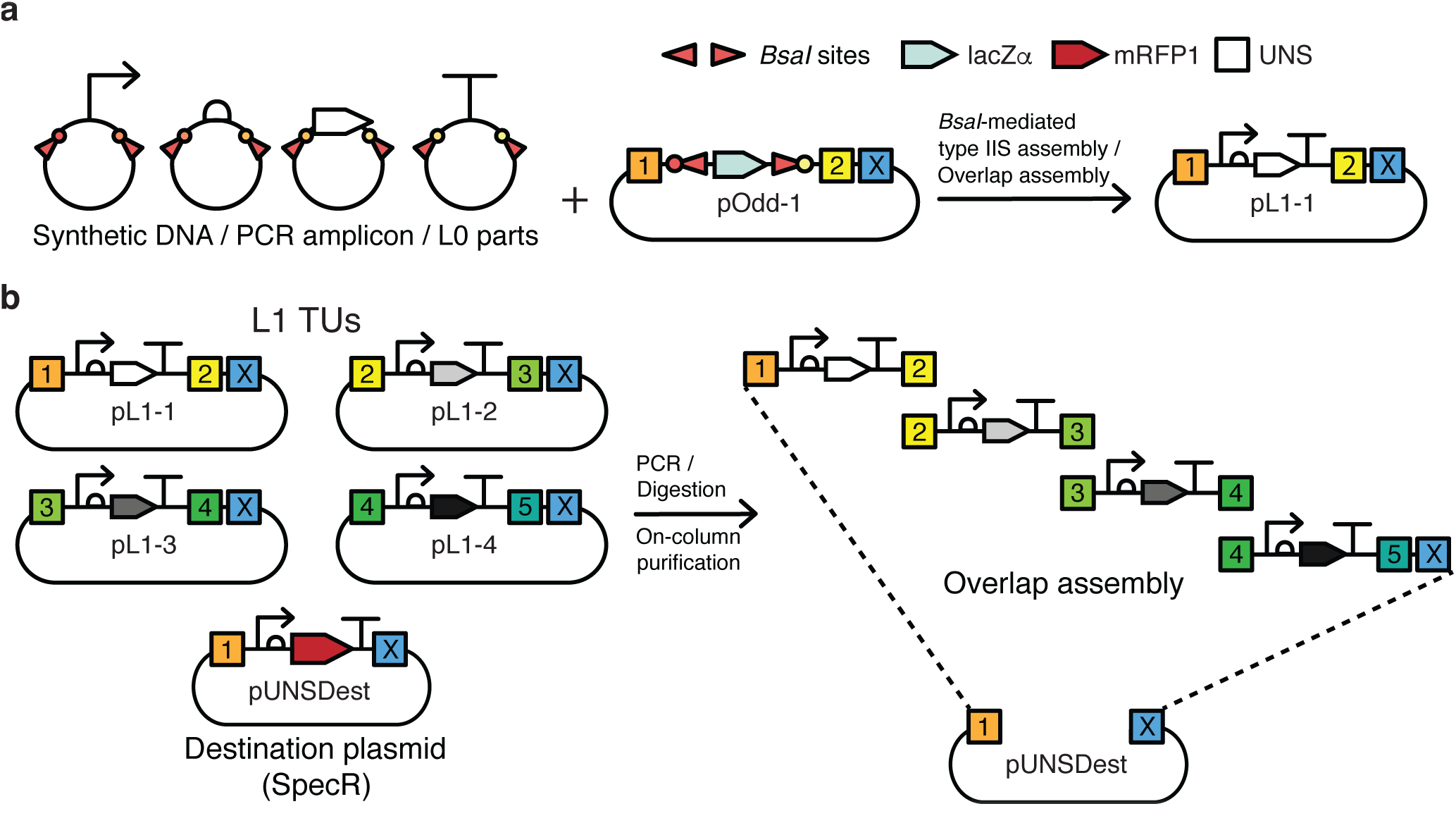
Loop overlap assembly. a, TU assembly for overlap assembly. UNS flanked TUs can be generated either by standard L0 BsaI-mediated Type IIS assembly or by overlap assembly methods using PCR-fragments or DNA synthesis. TUs produced by overlap assembly are only compatible with the overlap assembly pathway but do not require domestication. **b, Standardised overlap assembly**. Linear UNS flanked TUs are amplified by PCR or excised from plasmids by digestion by uncommon restriction enzymes. Linear UNS flanked TUs are then assembled to the destination plasmid pUNSDest by overlap assembly methods.

### Reliability of Loop assembly

To evaluate the technique’s reliability, we tested Loop assembly in different laboratories. We assembled over 200 plasmids using the Type IIS pathway for Levels 1-3 and obtained average assembly efficiency between 83 to 97 % depending on the level of assembly and complexity of constructs (Table 1, **Supplementary Text 3**). This was evaluated directly by means of restriction endonuclease digestion of the assembled plasmids. Further, we performed Illumina sequencing of 92 Level 2 and Level 3 assembled constructs to validate Loop assembly fidelity at the sequence level, to determine if the reaction had produced correct assemblies and if mutations had been introduced by our method. We found that 95.4 % of constructs assembled correctly with 98.8 % of overhang scars present at expected junctions. Overall, 99.8 % of nucleotides were correctly assembled, and the few incorrect constructs showed missing regions due to misassembly, rather than sequence errors *per se* (**Supplementary Table 3**).

### *In planta* activity of Loop plasmids

Loop vectors were derived from the pGreenII39 plant binary transformation vector. As in pGreenII, Loop plasmids contain elements for propagation in *Agrobacterium tumefaciens*, and are useful for *Agrobacterium*-mediated plant transformation. Loop assembly enables the construction of multiple-TU constructs from elemental L0 parts in as few as 2 cloning steps, providing a powerful tool for engineering plant gene expression. Simple constructs can be built with relative ease, and can be then composed into higher-level constructs. For example, combinations of fluorescent proteins with specific excitation and emission spectra, localization tags and specific promoters were assembled into useful reporters to highlight cellular features for their use in developmental studies. A Level 2 construct (pL2-1_TPL) containing a selectable marker (HygR), a mTurquoise2-N7 nuclear-localised reporter driven by a constitutive promoter (_*pro*_Mp*EF1* α^40,41^), a Venus-N7 nuclear-localised reporter driven by a tissue specific promoter (_*pro*_Mp*TPL*^42^) and a constitutive (_*pro*_Mp*EF1* α) eGFP-Lti6b membrane-localised marker was assembled from L0 parts (**Supplementary Table 4**) using Loop assembly and transformed into *Marchantia polymorpha* (Marchantia). Regenerated transformants were obtained and clonal propagules called *gemmae* were examined using confocal microscopy. All three fluorescent protein reporter genes were expressed and differentially localized in the transformed plants, and allowed visualization of distinct cellular and subcellular features across the tissue. The speed and flexibility of multiplex gene assembly provided by Loop assembly allows the facile generation of multispectral reporter constructs and their application to developmental studies (Fig. 5).

**Table 1.**
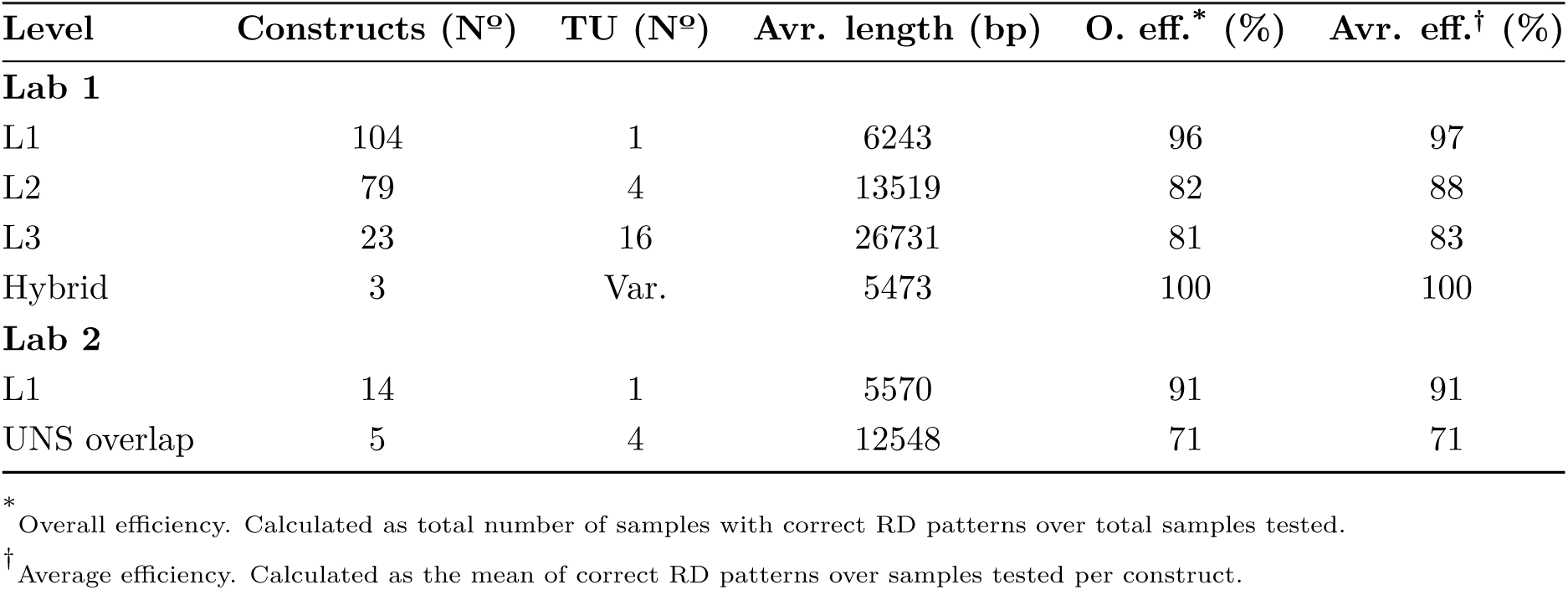
Loop assembly efficiency.

| Level | Constructs (N°) | TU (N°) | Avr. length (bp) | O. eff.* (%) | Avr. eff.† (%) |
| --- | --- | --- | --- | --- | --- |
| **Lab 1** |  |  |  |  |  |
| L1 | 104 | 1 | 6243 | 96 | 97 |
| L2 | 79 | 4 | 13519 | 82 | 88 |
| L3 | 23 | 16 | 26731 | 81 | 83 |
| Hybrid | 3 | Var. | 5473 | 100 | 100 |
| **Lab 2** |  |  |  |  |  |
| L1 | 14 | 1 | 5570 | 91 | 91 |
| UNS overlap | 5 | 4 | 12548 | 71 | 71 |

\* Overall efficiency. Calculated as total number of samples with correct RD patterns over total samples tested.
† Average efficiency. Calculated as the mean of correct RD patterns over samples tested per construct.

**Figure 5.**
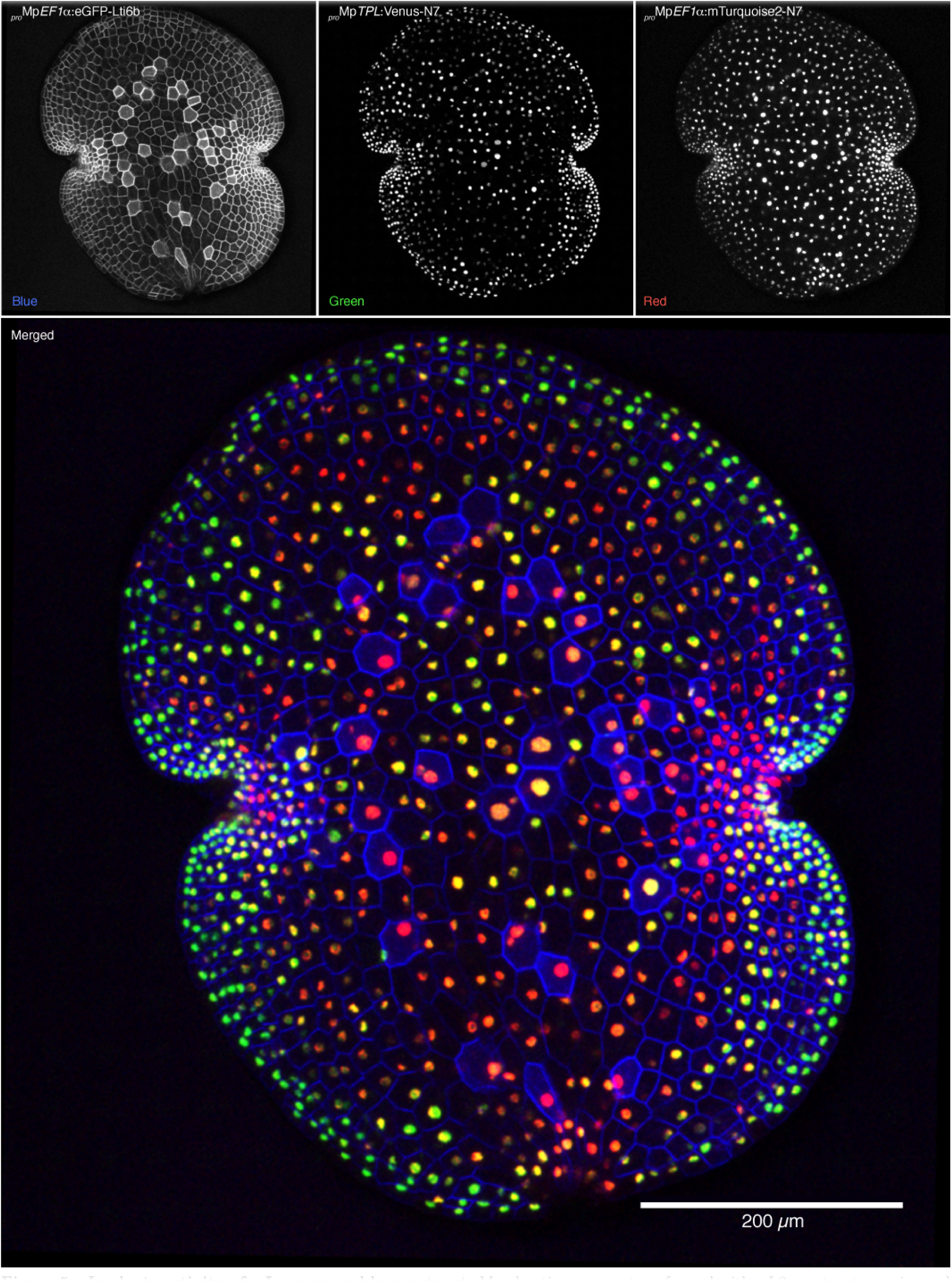
*In planta* activity of a Loop assembly construct. Marchantia *gemmae* transformed with a L2 construct were imaged through a Leica SP8 laser-scanning confocal microscope to assess expression of fluorescent markers. mTurquoise2-N7, Venus- N7 and eGFP-Lti6b were excited with appropriate wavelengths and fluorescence was captured in their respective emission windows in sequential scanning mode. Images shown are Z-stack maximum intensity projections.

In addition, 4 L1 TUs that had been constructed by Type IIS Loop assembly were built as a multi-TU destination plasmid using Gibson assembly. Transfected protoplasts showed expression of the engineered fluorescent reporters in their expected localizations (**Supplementary Fig. 4**), providing a fast and efficient system to evaluate functionality of Loop constructs. Maps of both plasmid constructs are provided in **Supplementary Fig. 5.**

### Loop assembly design automation

We have developed software tools to aid Loop assembly experiments. We developed *LoopDesigner*, a *Python*-based web application that facilitates (i) the sequence design and domestication of Level 0 DNA parts, (ii) generation of a Loop assembly parts database, and (iii) simulation of Loop assembly reactions and the resulting plasmid maps and sequences (Fig. 6). An input sequence is domesticated by identifying unwanted RE sites in a sequence and removing them by the introduction of synonymous mutations. Appropriate BsaI overhangs are added according to the rules of the common syntax for DNA parts. Loop assembly features a strictly ordered and recursive process, and assembly schemas were mapped and implemented using object-oriented software routines. Schemas were defined by choosing restriction enzymes and overhangs that establish the Type IIS Loop assembly type logic. See methods for detailed description of the *LoopDesigner* implementation. We invite readers to visit the *LoopDesigner* web tool available at loopdesigner.herokuapp.com (supported in Google Chrome) for exploring Loop assembly techniques. The source code of *LoopDesigner* is available at *GitHub* (https://github.com/HaseloffLab/LoopDB *LoopDesigner* branch), and provided under a MIT license.

**Figure 6.**
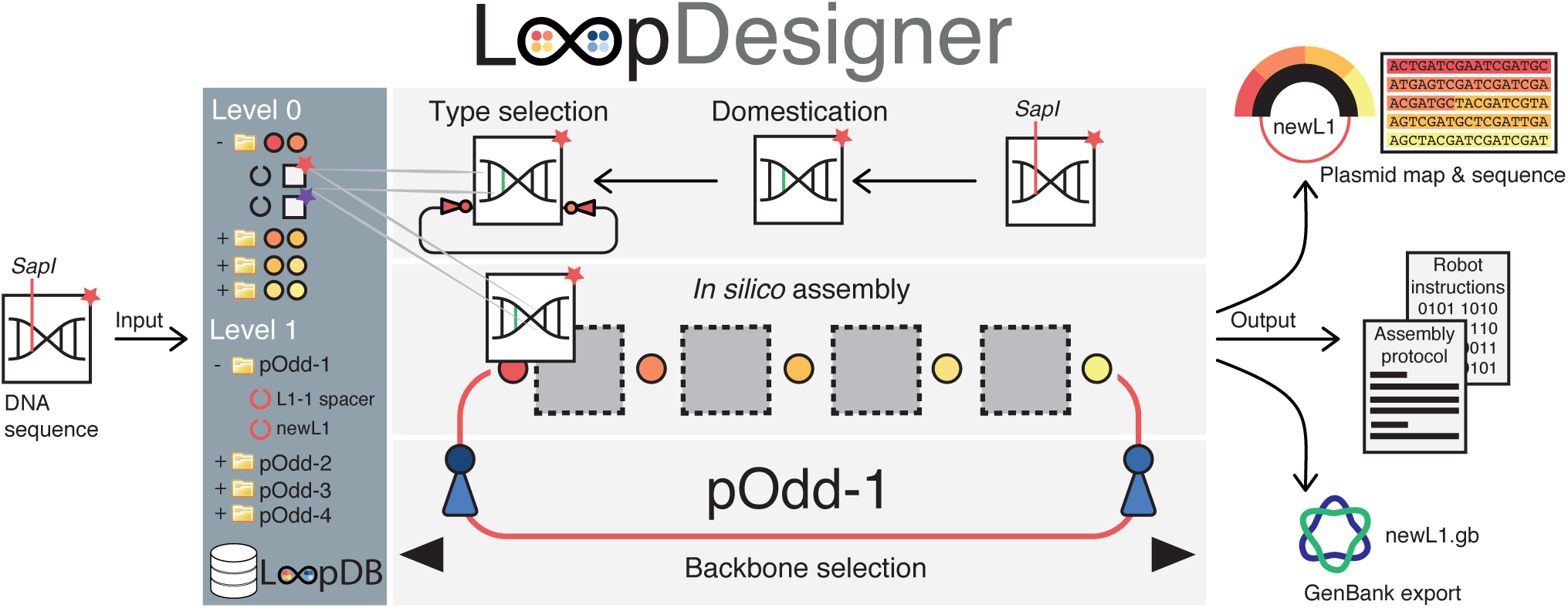
Design automation. A DNA sequence is submitted to *LoopDesigner*, which screens for BsaI and SapI sites and domesticates them to silent mutations where possible. A *part* type is specified for the assembly schema to save the *part* to the database library. To perform an *in silico* assembly, a receiver plasmid is selected which displays the compatible *parts* that can be placed in the current position of the assembly schema. As *parts* are included, the next compatible *parts* are displayed. When the assembly schema finds that the all *parts* required to complete the assembly are selected, the assembly simulation is performed. Then, *LoopDesigner* outputs the resulting plasmid map with its concurrent highlighted sequence and a protocol for Loop Type IIS reaction setup or export of GenBank sequence. Instructions to robots can be outputted if an API is provided with the required information (plasmid positions, ID mappings, robot functions) to produce the concurrent instruction file using *Python* scripting. The assembled *part* is then saved into the *part* library database for further assembly.

## Discussion

Loop assembly’s design is inspired by existing assembly methods such as GoldenBraid, Mo-Clo, and standardised Gibson assembly, but focused on the integration of these techniques into a general-purpose DNA assembly system. Loop assembly combines recursive use of two restriction enzymes and plasmid libraries, which together create a simple and versatile Type IIS assembly platform. Type IIS RE sites are employed in head-to-head configurations in two levels of assembly. The assembly schema eliminates the requirement for end-linkers, since the restriction sites for successive levels are integrated in receiver plasmids. Instead, Loop assembly requires all four ‘positions’ to be filled by either TUs or by spacers. Since introduction of a new RE imposes an additional constraint and requirement for domestication, we used SapI, a 7 bp recognition site RE, therefore diminishing the probability of finding its site fourfold, compared to an alternative RE with a 6 bp recognition site. The cloning system maintains the capacity for long sequence overlap assembly. Type IIS restriction sites are flanked by standardised UNS, enabling the use of Loop vectors with overlap assembly methods. We have demonstrated the high efficiency of the Loop assembly by generating a variety of constructs with different number of TUs, including the design and assembly of synthetic promoters. The ability to combine parts from different levels in a hybrid assembly is an intrinsic benefit of the recursive Loop assembly scheme.

The integration of different assembly techniques aids the generation of libraries of standardised DNA parts. A standard such as the common syntax enables the provision of tools and DNA parts for genetic engineering irrespective of the assembly method used. As Loop assembly integrates Type IIS and overlap assembly, it encourages the development of a community around a DNA construction system, yielding a growing collection of DNA parts and composites. The wide compatibility of Loop assembly facilitates proper curation and improvement of DNA part collections through collaboration, easier exchange and transfer of genetic modules between labs, and cross-validation. The ability to use either overlap or Type IIS assembly provides further flexibility in making DNA constructions where sequence alterations introduced by removal of illegal RE sites are not desirable (such as for experiments involving native genetic sequences), or when the assembly fails by one of the pathways.

Use and characterisation of the products of Loop assembly demonstrated that it is a robust and reliable DNA assembly system regardless of levels and types of parts. The high rate of successful assemblies, even in the absence of cPCR pre-selection, considerably decreases the effort and time required for DNA construction. Further, the system’s logical and technical simplicity enables rapid adoption by students and non-specialists, even for the generation of multiple TU constructs. This is due to the layers of abstraction provided by the use of a common syntax for DNA elements, the use of a simple scheme of plasmid vectors during assembly, common laboratory procedures and reagents, and streamlined protocols for the design and set up of Loop assembly reactions.

DNA fabrication is being propelled by laboratory automation platforms, chemical synthesis and design software. Although the falling costs of DNA synthesis suggest that DNA synthesis of transcriptional units or even chromosomes might eventually be time and cost-effective, synthetic biology requires the capacity for rapid, high-throughput and combinatorial assemblies. This is necessary for characterisation and troubleshooting of smaller DNA parts and circuits before compiling high-level devices and systems. Loop assembly provides all the intermediates in the process of hierarchical assembly, which can be repurposed or rearranged if required. In this respect, Type IIS assembly is particularly suited to automated assembly. Automated design and liquid handling platforms for fabrication DNA constructs have already been adopted by some^32^ and the technologies are rapidly expanding: at the high end of the market, platforms such as the Echo (Labcyte) are enabling miniaturisation, and increasing throughput^43^, while low-cost platforms such as the OT-One S (OpenTrons) are aiming to make automated pipetting possible in every laboratory.

To enable the rapid and automated design of constructs that can be assembled on high-throughput automated platforms we developed *LoopDesigner*, a software framework that provides an interface between digital design and experimentation which can be easily used to tether robotic platforms with Loop assembly. We have demonstrated the power of the *LoopDesigner* by implementing a simple web tool where users can design assembly strategy and run virtual reactions before stepping into the lab. The *LoopDesigner* framework allows definition of Loop assembly schemas of arbitrary complexity with any number of levels and plasmids per level, as well as with any possible restriction enzymes and overhangs, as long as the defined parts have matching over-hangs. Furthermore, *LoopDesigner* generalises the concept of the assembly, so that the assembly schema presented in this paper become a single instance of many possible implementations of the Loop assembly, allowing for the exploration of novel ways of assembling DNA parts through Type IIS strategies.

DNA construction has traditionally coupled the process of plasmid assembly and concurrent use of plasmids in model organisms. Loop assembly provides the sufficient throughput and versatility for working with general-purpose backbones, in which users can add specific traits e.g. parts for transfection. Vectors could be decoupled from specific uses by modularising replication origins and selection markers as basic DNA parts and introducing host-specific elements during the assembly process. This would provide higher flexibility during design, and allow switching selection markers when super-transformation is required, for instance. Such approaches would make the DNA fabrication process host-agnostic, promoting the development of universal DNA assembly systems using standards such as the common syntax, which would provide unprecedented exchange of DNA components within the biological sciences.

Until recently, the majority of materials for research were exchanged under a Uniform Biological Material Transfer Agreement (UB-MTA). This is a bilateral legal agreement that, in its standard form, does not allow redistribution, exchange or use with those outside of educational and research organisations such as universities and government research institutes. At the same time, in scientific publishing and in software, there is a trend toward openness to facilitate collaboration and translation of basic research. An excellent example of how the open source philosophy has powered and enabled innovation is exemplified by the software development field by community-based coding projects such as the ones hosted by *Github* (https://github.com). *Git* was originally developed for the purpose of coordination and collaboration of distributed software development, and nowadays most collaborative projects (both in the public and the private sector) use *Git* as an underlying framework. It is unlikely that we will see such thriving success in DNA engineering and synthetic biology, unless new forms of unrestricted DNA sharing and assembly are established under more open frameworks such as the OpenMTA. We support the adoption of an open-source inspired L0 elemental part exchange by providing Loop assembly for the higher-level construction of these L0 components under an OpenMTA framework. Work on establishing an OpenMTA is underway to ensure access to the Loop assembly system remains democratic and unhindered for both the public and the private sector.

## Methods

### Construction of Loop assembly backbones

Loop assembly vectors were constructed using Gibson assembly (Gibson, *et al*. 2009^3^). Starting from a pGreenII vector (Hellens, et al. 2000^39^), several changes were made to obtain a basic plasmid backbone for the Loop assembly vectors: BsaI and SapI sites were removed from the plasmid and recoded using silent mutations. The pGreenII ColEI origin of replication was mutated to the low copy number pBR322 origin of replication by mutating 2 nucleotides to reduce issues with DNA replication of large constructs in bacteria. A region extending from the T-DNA left border to the hygromycin resistance gene cassette was replaced with the sequence of the pET15 vector^44^ from the nptII nosT terminator to the UASGAL4 promoter (bases 2851- 3527). A spectinomycin resistance was cloned to replace of the nptI cassette to provide a microbial selection marker for the pEven plasmids. UNS were cloned into the kanamycin and spectinomycin version of vector backbones after the 3’ end of the pET15 vector sequence and the RB. Finally, the Loop restriction enzyme sites (BsaI and SapI), overhangs and the lacZα cassette were cloned in between the UNS, yielding the pOdd and pEven vectors described in Fig. 2. L0 plasmids used for Loop Type IIS assembly were assembled using Gibson assembly into a modified pUDP2 (BBa_P10500) plasmid, which contained a 20 bp random sequence (5’-TAGCCGGTCGAGTGATACACTGAAGTCTC-3’) downstream of the 3’ convergent BsaI site and before the BioBrick suffix, to provide non-homologous flanking regions for correct orientation during overlap assembly.

### Plasmids and construct design

L0 parts used for DNA construction are described in **Supplementary Table 4,** their sequences included in **Supplementary File 1** and are available through Addgene. Plasmid maps for resulting multigene assemblies are included in **Supplementary Fig. 5** and sequences included in **Supplementary File 1.**

The design of the constructs was performed using *LoopDesigner* software, installed on a local machine. The software was configured to use Loop assembly backbones together with BsaI and SapI RE, as well as A-B and α-ω overhangs. In addition, definition of 12 L0 part types were added to the software, based on the overhangs specified by the common syntax. The sequences of the L0 parts were added to the *LoopDesigner* database, assigning one of the defined part types, and assembled consequently into Level 1 and Level 2 constructs *in silico.* The concentration of L0 parts and Level 1 constructs were adjusted to those suggested by the LoopDesigner for 10 μL reactions.

### Loop Type IIS assembly protocol

The Loop Type IIS assembly protocol was adapted from Patron, 201628. 15 fmol of each part to be assembled were mixed with 7.5 fmol of the acceptor plasmid in a final volume of 5 μL with dH_2_0 (**Supplementary Table 5**). The reaction mix containing 3 μL of dH_2_0, 1 μL of T4 DNA ligase buffer 10x (NEB cat. B0202), 0.5 uL of 1 mg/mL purified bovine serum albumin (1:20 dilution in dH_2_0 of BSA, Molecular Biology Grade 20 mg/mL, NEB cat. B9000), 0.25 μL of T4 DNA ligase at 400 U/μL (NEB cat. M0202) and 0.25 μL of corresponding restriction enzyme at 10 U/μL (BsaI NEB cat. R0535 or SapI NEB cat. R0569) was prepared on ice. 5 μL of the reaction mix was combined with the 5 μL of DNA mix for a reaction volume of 10 μL (**Supplementary Table 6)** by pipetting and incubated in a thermocycler using the program described in **Supplementary Table 7.** For SapI reactions, T4 DNA ligase buffer was replaced by CutSmart buffer (NEB cat. B7204S) supplemented with 1 mM ATP. 1 μL of the reaction mix was added to 50 μL of chemically competent TOP10 cells (ThermoFisher cat. C4040100) and following incubation at 42°C for 30 seconds, samples were left on ice for 5 minutes, 250 μL of SOC media was added and cells incubated at 37 °C for 1 hour. Finally, 5 μL of 25 mg/mL of X-Gal (Sigma-Aldrich cat. B4252) dissolved in DMSO, was added and the cells were plated onto selective LB-agar plates. Assembly reactions were also automated. The assembly reactions were identical except scaled down to a total volume of 1 μL. Reactions were set up on a LabCyte Echo in 384 well plates and incubated on a thermal cycling machine using the same conditions as described above. Reactions were transformed into 4 μL competent XL10-Gold^®^ Ultracompetent Cells (Agilent Technologies, Santa Clara, CA, USA) and plated onto eight-well selective LB-agar plates. Colonies were picked for growth in 1 mL of media in 96-well plates on a Hamilton STARplus^®^ platform.

### Standardised PCR of transcriptional units

PCR conditions used with UNS primers were 60 ºC, 35 cycles using Phusion High-Fidelity DNA polymerase (ThermoFisher cat. F-530) in 50 μL reactions, according to manufacturer’s instructions. Template was added to a final concentration of 20 pg/μL. DNA fragments were visualized using SYBR Safe DNA Gel Stain (Ter-moFisher cat. S33102) on a blue LED transilluminator (IORodeo). DNA purification was performed using NucleoSpin Gel and PCR Clean-up purification kit (Macherey-Nagel, cat. 740609.250). UNS primers used in TUs amplification are listed in **Supplementary Table 2**.

### Validation by sequencing

The sequences of assembled plasmids were verified by complete sequencing using 150 base pair paired-end reads on an Illumina MiSeq platform. Libraries were prepared using the Nextera XT DNA Library Prep Kit (Illumina cat. FC-131-1096), with the protocol modified a one in four dilution. Reads were filtered and trimmed for low-quality bases and mapped to plasmids using the ‘map to reference tool’ from the Geneious 8.1.8 software (http://www.geneious.com, Kearse *et al*., 2012^45^), with standard parameters. Sequence fidelity was determined manually.

### *Agrobacterium*-mediated Marchantia transformation

*Agrobacterium*-mediated transformation was carried out as described previously (Ishizaki, et al. 2008^46^), with the following exceptions: half of an archegonia-bearing sporangium (spore-head) was used for each transformation. Dried spore-heads were crushed in a 50 mL Falcon tube with a 15 mL Falcon tube and resuspended in 1 mL of water per spore-head. Resuspended spores were filtered through a 40 μm mesh (Corning cat. 352340) and 1 mL of suspension was aliquoted into a 1.5 mL Eppendorf tube and centrifuged at 13,000xg for 1 min at room temperature. Supernatant was discarded and spores were resuspended in 1 mL of sterilisation solution (1 Milton mini-sterilising tablet (Milton Pharmaceutical UK Company, active ingredient: Sodium dichloroisocyanurate CAS: 2893-78-9: 19.5% w/w) dissolved in 25 mL of sterile water, and incubated at room temperature for 20 min at 150 RPM on rotating shaker. Samples were centrifuged at 13,000xg for 1 min, washed once with sterile water and resuspended in 100 μL of sterile water per spore-head used. One hundred μL of sterilised spores were inoculated onto half strength Gamborg’s B5 1 % (w/v) agar plates and grown under constant fluorescent lighting (50-60 mol photons/m^2^s) upside down for 5 days until co-cultivation. Sporelings were co-cultivated with previously transformed and induced *Agrobacterium* GV2260 transformed with the pSoup plasmid (Hellens, et al. 2000^39^) in 250 mL flasks containing 25 mL of half strength Gamborg’s B5 media supplemented with 5 % (w/v) sucrose, 0.1 % (w/v) N-Z Amine A (Sigma cat. C7290), 0.03 % (w/v) L-glutamine (Sigma cat. G8540) and 100 μM acetosyringone (Sigma-Aldrich cat. D134406) for 36 h, until washed and plated onto selective media.

### Laser-scanning confocal microscopy

A microscope slide was fitted with a 65 μL Gene Frame (ThermoFisher cat. AB0577) and 65 μL of dH20 was placed in the center. Marchantia *gemmae* were carefully deposited on the drop of dH20 using a small inoculation loop and a thin coverslip was attached to the Gene Frame. Slides were used for confocal microscopy on a Leica TCS SP8 confocal microscope platform equipped with a white-light laser (WLL) device. Imaging was conducted using a Leica HC PL APO 20x CS2 air objective with a sequential scanning mode with laser wavelengths of 405 nm, 488 nm and 515 nm, capturing emitted fluorescence at 450-482 nm, 492-512 nm and 520-550 nm windows in each sequential scan, respectively. Z-stacks were collected every 5 μm for the complete volume range and maximum intensity projections were processed using ImageJ software. Fluorescence bleedthrough from the blue pseudocoloured channel (membrane localized eGFP) into the green pseudo-coloured channel (nuclear localized Venus) was eliminated using custom *Python* scripts which subtracted a 20% of the value of pixels present in the blue channel to the green channel. Images were edited to scale the pixel intensity to the full 8-bit range and a merged channel was processed.

### Transient expression in Arabidopsis mesophyll protoplasts

Well-expanded leaves from 3-4 weeks old Arabidopsis plants (Columbia-0) were used for protoplast transfection. Plants were grown at 22ºC, low light (75 μmol/m^2^s) and short photoperiod (12 h light/12 h dark) conditions. Protoplasts were isolated and PEG-transfected according to Yoo*, et al.* 2007^47^. For transfection, 6 μL of Loop L2 plasmids (2 μg/μL) isolated by NucleoBond Xtra Midi/Maxi purification kit (Macherey-Nagel cat. 740410.50), were used. Transfected protoplasts were incubated for 12 hours in light and then visualized by epi-fluorescent microscopy in a Neubauer chamber (Hirschmann).

### Epifluorescence microscopy

Transfected protoplasts were visualized using a Nikon Ni microscope equipped with a 49021 ET - EBFP2/Coumarin/Attenuated DAPI filter cube (ex: 405/20 nm, dichroic: 425 nm, em: 460/50 nm), 96227 AT-EYFP filter cube (ex: 495/20 nm, dichroic: 515 nm, em: 540/30 nm), 96223 AT-ECFP/C filter cube (ex: 495/20 nm, dichroic: 515 nm, em: 540/30 nm) and a 96312 G-2E/C filter cube (ex: 540/20 nm, dichroic: 565 nm, em: 620/60 nm).

### Bacterial transformation of Type IIS reactions

A volume of 50 μL of chemically competent TOP10 cells (ThermoFisher cat. C404010) was thawed on ice for 10 min and mixed with 1 μL of a Loop Type IIS assembly reaction. Samples were subjected to heat-shock for 30 seconds and left on ice for 5 minutes, where 250 μL of SOC media was added. Samples were incubated at 37 ºC for 1 hour and 50 μL of sample was mixed with 5 μL of 25 mg/mL of X-Gal (Sigma-Aldrich cat. B4252) dissolved in DMSO, and inoculated onto selective LB-agar plates.

### LoopDesigner

In order to implement an object oriented model for Loop assembly, we built a PartsDB library (https://github.com/HaseloffLab/PartsDB) to define several interlinked classes, each of which is associated with a table in a relational SQL database. The structure of *LoopDesigner* is built around a *Part* class, which either represents an ordered collection of child parts it is assembled from, or a DNA sequence in case of L0 parts. In this way we ensured that the actual DNA sequence is only stored once, while the sequences of L1 and higher parts are constructed on demand from the relational links. In addition, each *Part* is associated with one of the *Backbone* instances, which together with a *Part* sequence represents a complete Loop assembly plasmid. Every instance of a *Backbone* class is a combination of a *Base Sequence* and a donor Restriction Enzyme Site, e.g. pOdd 1-4 and pEven 1-4 are *Backbone* instances in the schema described in this paper. *Base Sequence* represents a type of a receiver plasmid, e.g. pOdd and pEven, and is composed of a DNA sequence of the plasmid and an instance of a receiver *Restriction Enzyme Site*. Finally, *Restriction Enzyme Site* class is composed of a *Restriction Enzyme* instance, which stores restriction enzyme recognition sequence, and a pair of overhang sequences, which can be either receiver or donor.

## Acknowledgements

We would like to acknowledge Diego Orzaez from Instituto de Biología Molecular y Celular de Plantas for helpful suggestions and advice regarding SapI and Type IIS restriction enzymes. Oleg Raitskin for advice on Type IIS assembly protocols, H. Ghareeb from Göttingen University, T. W. J. Gadella, and J. Goedhart from University of Amsterdam for providing mTurquoise2 DNA, Jen Sheen from Harvard University for providing TCS promoter DNA, Teva Vernaux from of Ecole Normale Supérieure de Lyon for Venus N7 DNA, and Jeff Lichtman from Harvard University for mTagBFP2 DNA. Susana Sauret-Gueto, Eftychis Frangedakis and Marta Tomaselli from the University of Cambridge for troubleshooting SapI reactions. We thank Linda Kahl and Drew Endy, BioBricks Foundation for discussions about implementation of the OpenMTA.

**Correspondence** Correspondence and requests for materials should be addressed to F.F. or J.H.

